# Are veterinary clinical manuscripts published more slowly than medical or scientific counterparts? A comparative observational study

**DOI:** 10.1101/456327

**Authors:** Mark Rishniw, Maurice E. white

**Author notes:** Corresponding author. M.R., E-mail address (M.R).

## Abstract

**BACKGROUND:** Publication speeds of clinically relevant veterinary journals have not been evaluated.

**METHODS:** We compared 23 prominent veterinary journals to 11 comparable medical and 4 high-impact science journals and examined select factors that might affect these speeds. Submission date, acceptance date and first online publication date were recorded for 50 sequentially identified research articles from each journal that had been published immediately prior to April 26, 2018. Intervals from submission to acceptance, acceptance to publication, and submission to publication were calculated in days for each article. Data were compared visually across all journals, and statistically by field (veterinary, medical, high-impact), by impact factor and by publisher (commercial vs society) to identify trends or differences in publication intervals.

**RESULTS:** When assessed by field, intervals from submission to acceptance (p=0.18), acceptance to publication (p=0.75) and submission to publication (p=0.13) did not differ. Individual journals varied slightly in intervals from submission to acceptance, but varied markedly in intervals from acceptance to publication. Three journals had median intervals from acceptance to publication exceeding 135 days and two exceeding 500 days. Three journals had median intervals from submission to publication exceeding 550 days. Neither impact factor nor publication model affected any intervals. Intervals from submission to acceptance and acceptance to publication were positively associated with overall interval from submission to publication (rho=0.7, P<0.0001 for both associations).

**CONCLUSIONS:** Intervals from submission to acceptance for veterinary journals are like those for medical and high-impact journals, suggesting that the review process is similar across fields. However, several veterinary journals have intervals from acceptance to publication approaching 18 months.

---

THE NUMBER OF PEER REVIEWED veterinary, medical and scientific articles over the last 30 years has increased dramatically, from approximately 200,000 in1973 to 1.2 million in 2013.^1^ The development of the internet and computer software has facilitated the acquisition, analysis, preparation and presentation of data, and the submission process to journals, allowing this rapid expansion of published material.

Clinicians and researchers depend on timely dissemination of information.^2^ Rapid dissemination allows patient management strategies to advance and promotes further research. Furthermore, it reduces unnecessary duplication or wasted research efforts. Indeed, critics have argued that investigators conducting publicly funded clinical trials have a responsibility to publish results rapidly.^3^ Rapid publication also assists young investigators with promotion and securing employment. However, perceptions exist amongst clinicians ^4^, and both authors and journal editors have expressed concerns that publication speed is unreasonably slow, and have proposed potential solutions.^5–7^

Various investigators have examined publication speed of journals within biomedical and other scientific fields^8–14^ Investigators have variously identified or excluded factors as contributing to publication speed, including impact factor, type of study, journal type (open-access vs “reader pays”, society-published vs. publishing company). ^7^^,^^11^^,^^12^^,^^14^^,^^15^ Most studies examine two components of publication speed: interval from submission to acceptance, and interval from acceptance to either first appearance online or in print. To our knowledge, no studies have examined these components for publication of veterinary clinical journals.

Therefore, we examined the intervals from submission to acceptance, acceptance to publication and submission to publication of prominent clinical veterinary journals and compared them to that of select comparable medical journals and a small cohort of general scientific journals with high impact factors. We also examined whether impact factor or journal publisher (commercial vs society) affected publication speed across all journals in our study.

## METHODS

### JOURNAL SELECTION

We obtained a list of 234 veterinary journals, ranked by impact factor (from highest to lowest), and from this list, we selected English-language veterinary journals that we considered relevant to most veterinary clinicians. We based “relevance” on criteria used by one of the authors (White) who examines and selects articles for CONSULTANT (online clinical reference database). This author had previously shown that a relatively small subset of journals provides most of the content for CONSULTANT,^16^ consistent with Bradford's law.^17^ This produced a final list of 31 veterinary journals. We then selected 15 comparable representative clinical medical journals and 4 general scientific journals with very high impact factors from which we hoped to extract data. Our process for selecting clinical medical journals was based on journal reputation or familiarity (e.g. New England Journal of Medicine, Journal of the American Medical Association, Lancet) and comparability to veterinary clinical journals that we had included in our initial list. We identified high-impact general science journals by examining an online list of journals ranked by impact factor.

The sample was a convenience sample and did not include any formal randomization or selection criteria other than those described above.

### DATA EXTRACTION

We examined each of the journal websites for the following information, which was provided as a metric with each article: date of initial manuscript submission, date of final acceptance and date of first online publication.

In journals with complete data, we systematically examined the 50 most recent research articles in each journal published before April 26, 2018, by examining the table of contents for each issue and identifying the relevant publication metrics. Once data from 50 articles had been recorded, we ceased data collection for that journal. Therefore, depending on the frequency of publication, and the number of articles per issue, the period over which we collected data differed by journal, but did not exceed 6 months for any journal. We manually recorded the submission date, acceptance date and first online publication date, usually reported on the manuscript website as “first record online” or “published online”. From these dates, we calculated, in days, the interval from submission to acceptance, the interval from acceptance to publication and the interval from submission to publication.

Four medical journals and 9 veterinary journals did not provide these data with the articles; further, 3 medical journals, one high-impact journal and one veterinary journal provided partial information (Table 1). Because one of the authors was a scientific editor with the veterinary journal providing partial information (The Veterinary Journal), he extracted the missing information (submission date) from the manuscript histories. Furthermore, because the Journal of the American Veterinary Medical Association is considered a flagship journal in the United States for the veterinary community, and the online version is published simultaneously with the print edition, one of the authors contacted the corresponding authors of the 70 most recent research articles (as of April 26, 2018) by email and asked them to provide the submission dates and acceptance dates for their manuscripts. We analyzed data from the first 50 respondents to our request. Finally, for the one high-impact journal with partial information (Science), we collected data from “early view on line” manuscripts, rather than those published in the paper issues (where only the submission and acceptance dates were available). Therefore, we ultimately evaluated 23 veterinary journals (all with complete information), 4 high impact journals (all with complete information) and 12 medical journals (three with partial information). The three journals with partial information were excluded from the analyses that would have required the missing data.

**Table 1.**
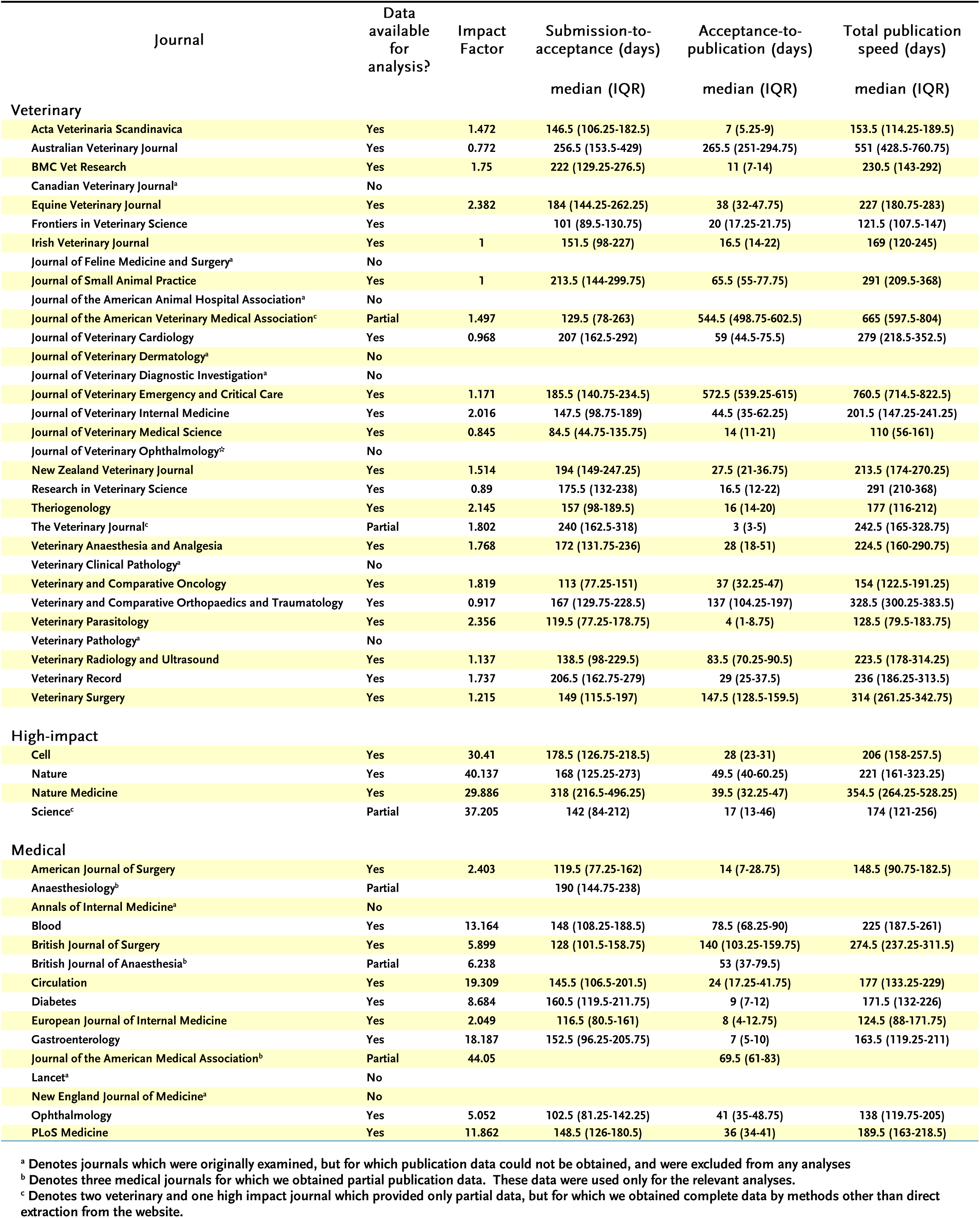
Impact factors, submission-to-acceptance times, acceptance-to-publication times and total publication speed for veterinary, medical and high-impact journals analysed in this study

We excluded special issues, review articles, case reports, commentaries, opinion pieces and editorials. For journals with partial data consisting of at least two dates, we recorded those dates, and restricted analyses of those journals accordingly. We recorded impact factors for each journal provided either by a Google search or from the journal websites, as available on April 22, 2018. We categorized each journal as either “commercially published” if the journal was published by a commercial publishing company (e.g. Elsevier, Wiley, Springer-Link etc.), or “society published” if the publishing group was the journal organization (e.g. American Veterinary Medical Association, Canadian Veterinary Association, American Diabetes Association, etc.).

### STATISTICAL ANALYSIS

To examine whether intervals from submission to acceptance or acceptance to publication differed by field (i.e., veterinary, medical or high-impact), we compared intervals by Kruskal-Wallis tests.

We examined the associations between impact factor and the interval from submission to acceptance, acceptance to publication and submission to publication using Spearman's Rank Correlation. We further examined whether the type of publisher (i.e., commercial or society) affected the interval from submission to acceptance, and acceptance to publication by rank sum tests. Finally, we examined whether the interval from submission to acceptance, or acceptance to publication was associated with total time from submission to publication by Spearman's Rank Correlation.

We plotted the publication intervals (submission to acceptance, acceptance to publication, and submission to publication) using box and whisker plots. We did not compare these intervals between individual journals, but only examined the data visually.

## RESULTS

### JOURNALS HAVE SIMILAR INTERVALS FROM SUBMISSION TO ACCEPTANCE

Data were available for all journals except the British Journal of Anaesthesia and the Journal of the American Medical Association (Table 1). Figure 1 shows the intervals from submission to acceptance for all journals. Intervals from submission to acceptance did not differ by field (veterinary: 167 days [IQR: 139 to 207, n=23]; high-impact: 173 days [IQR: 149 to 283, n=4]; medical: 147 days [IQR: 119 to 155, n=10]; p=0.13; Figure 2). Nature Medicine had the longest median interval (318 days; IQR: 217 to 496).

**Figure 1.**
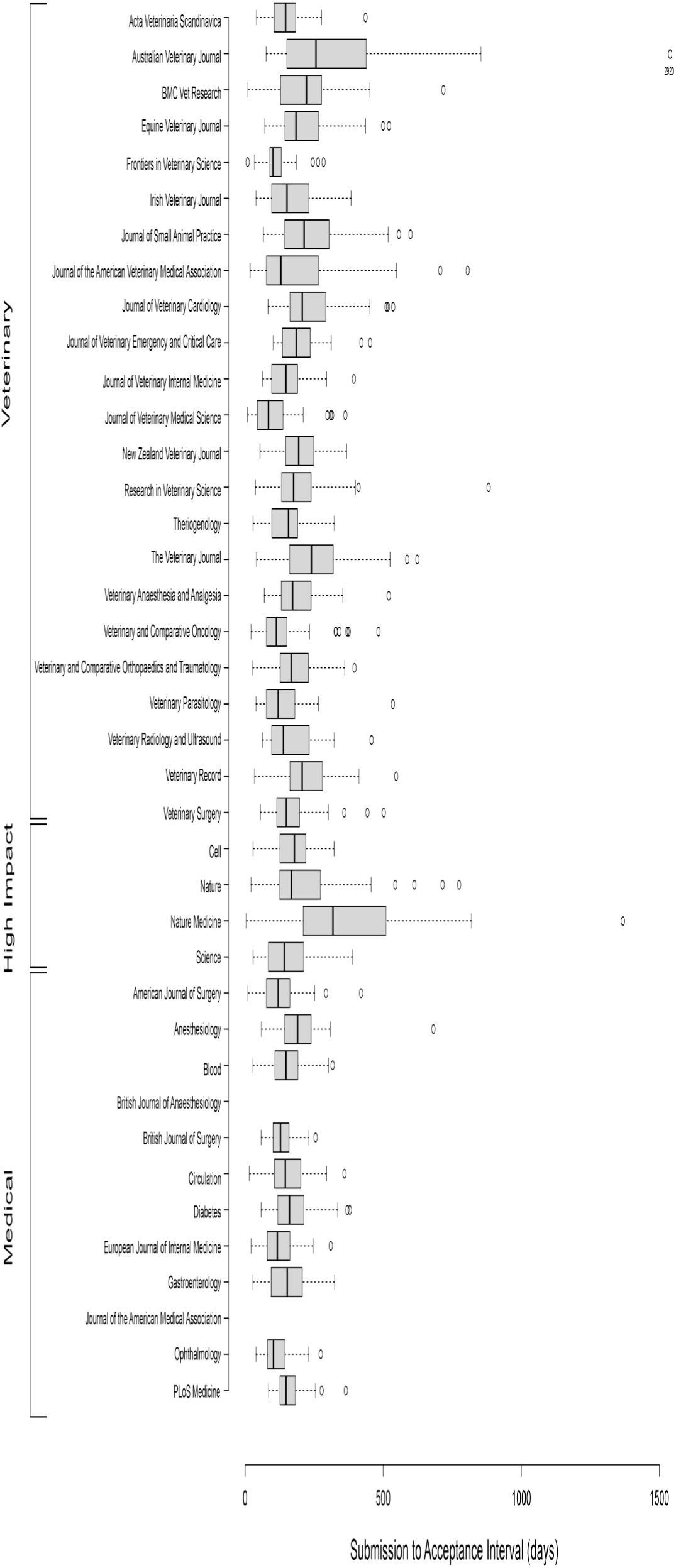
Box-and-whisker plots of the intervals from submission to acceptance of 50 articles for select veterinary, medical and high-impact journals. Whiskers denote the 5th and 95th percentiles. Circles denote articles outside the 5th and 95th percentiles.

**Figure 2.**
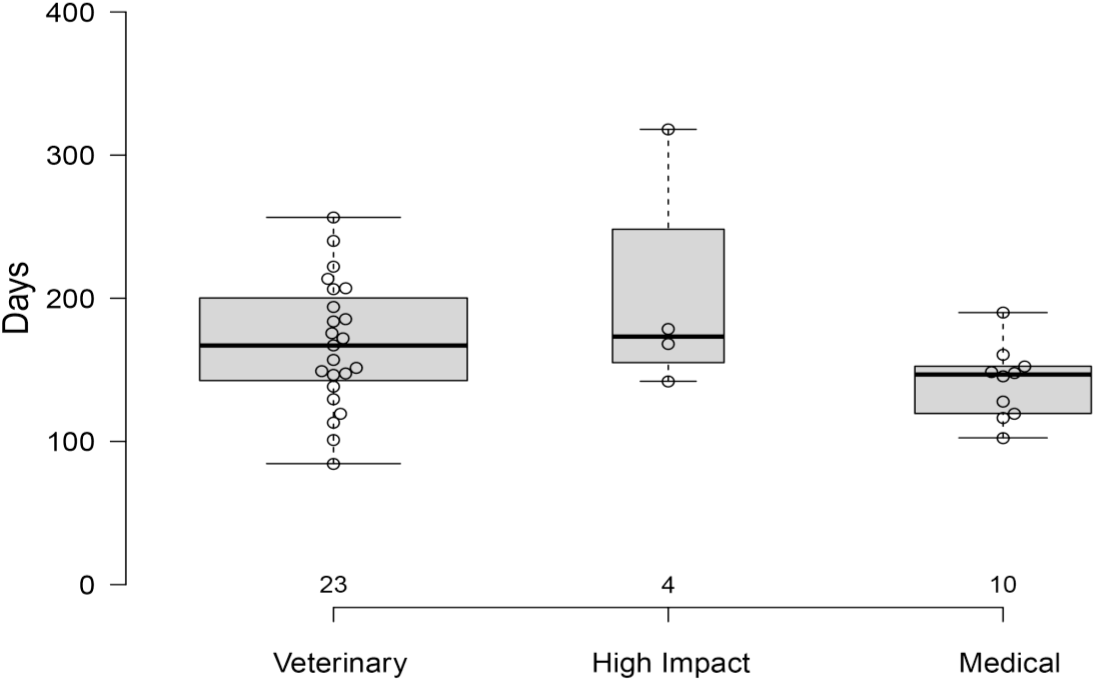
Box-and-whisker plots of median intervals from submission to acceptance in journals grouped by field (veterinary, medical, high impact). Groups did not differ from each other. Whiskers denote values within 1.5x the interquartile range. Circles denote individual values.

Society-published journals (median: 144 days [IQR:116 to 148, n=8]) accepted articles faster than commercially published journals (median: 168 days [IQR:147 to 203, n=27]) (p=0.01). The interval from submission to acceptance was/was not associated with impact factor (rho= -0.164, p=0.34, Figure 3A). However, the interval from submission to acceptance was associated with the total interval from submission to publication (rho=0.7, p<0.001, Figure 4A).

**Figure 3.**
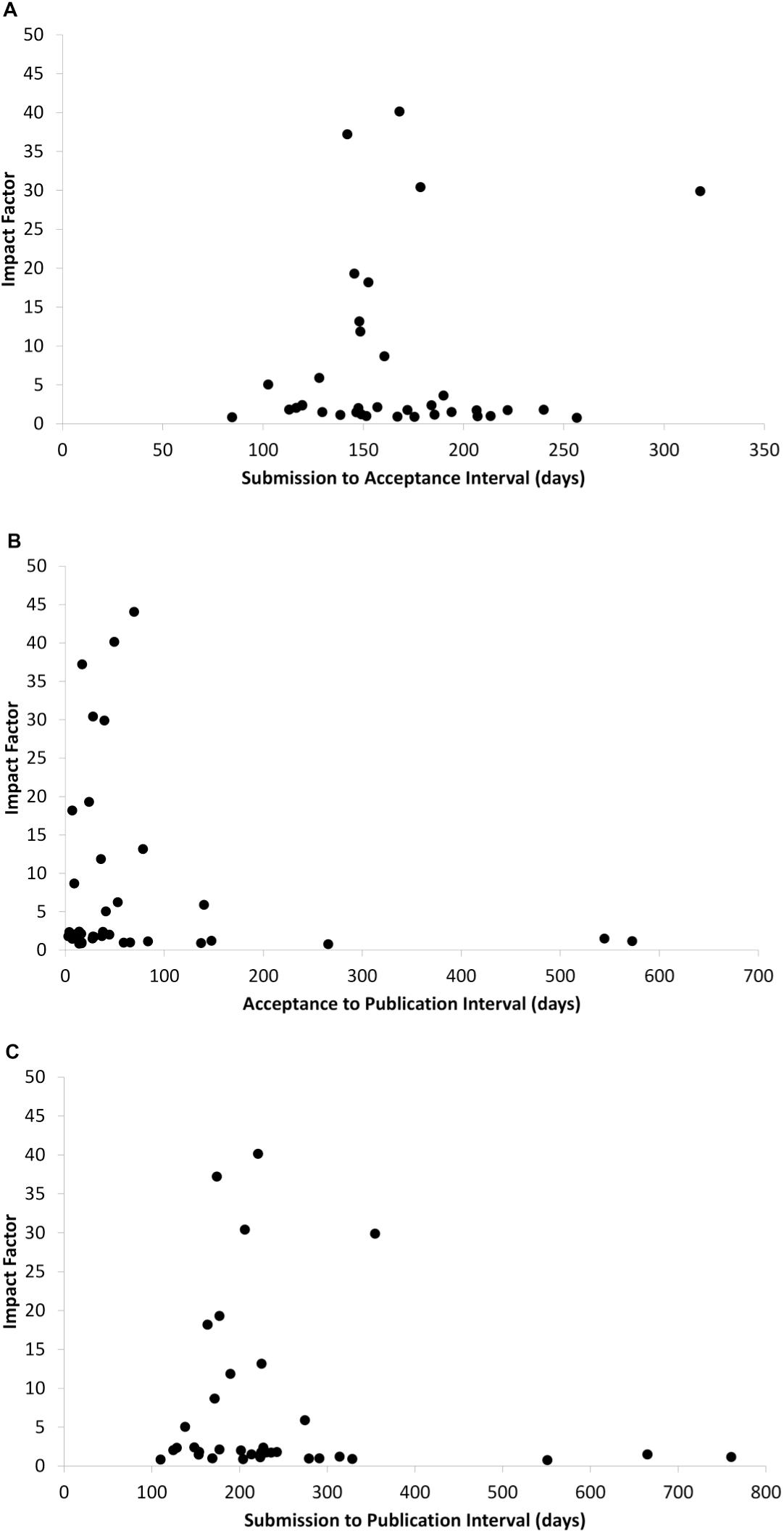
Scatter plots of the relationship between impact factors for each journal and the median intervals from (A) submission to acceptance, (B) acceptance to first publication online and (C) submission to first publication online of 50 articles for select veterinary, medical and high-impact journals.

**Figure 4.**
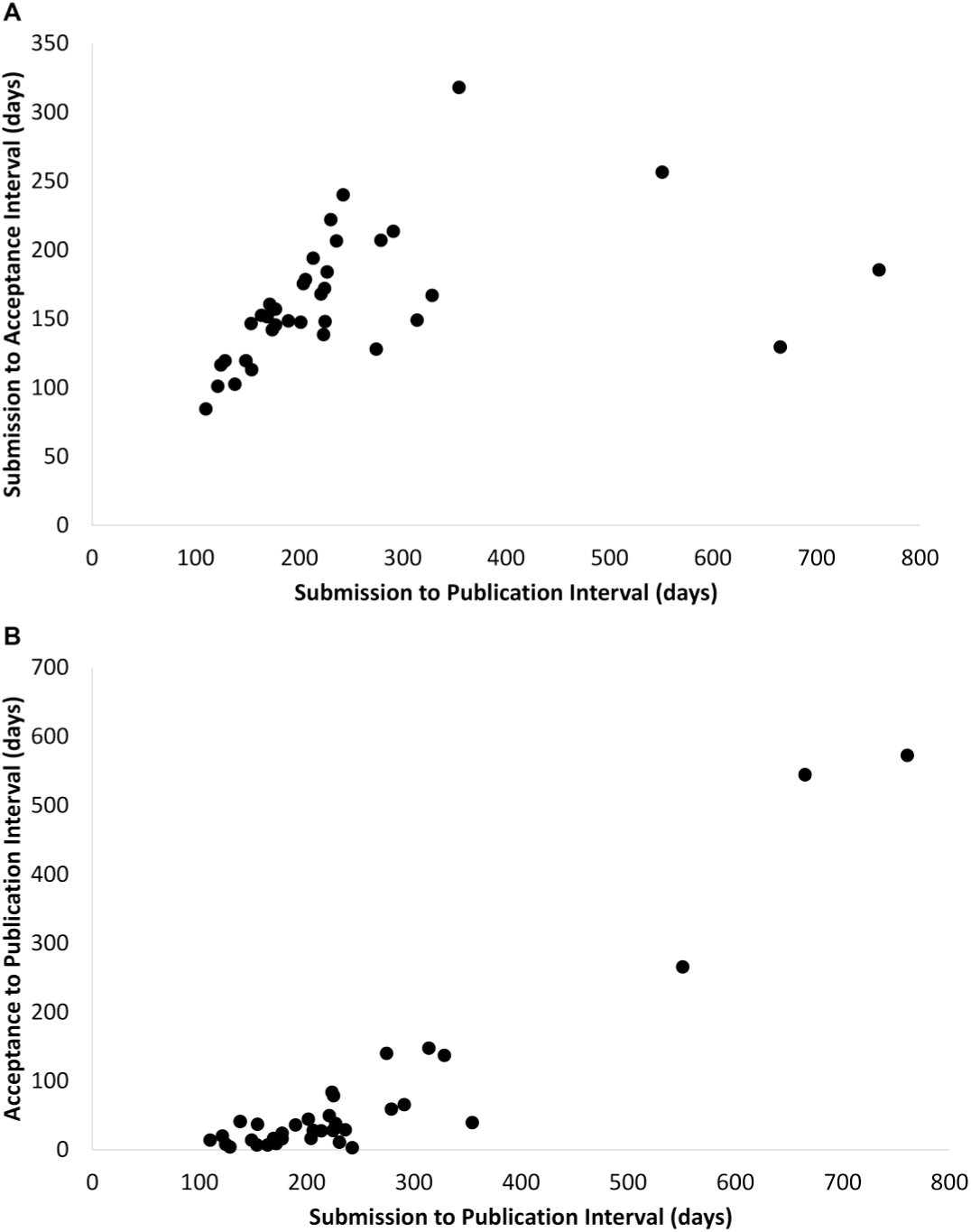
Scatter plots of the relationship between the median interval from submission to first publication online of 50 articles for select veterinary, medical and high-impact journals and the median intervals from (A) submission to acceptance and (B) acceptance to first publication online.

### SOME VETERINARY JOURNALS HAVE LONG INTERVALS FROM ACCEPTANCE TO PUBLICATION

Figure 5 shows the intervals from acceptance to publication for all journals (Table 1). Intervals from acceptance to publication did not differ by field (veterinary: 29 days [IQR: 16 to 79, n=23]; high-impact: 34 days [IQR: 22 to 45, n=4]; medical: 36 days [IQR: 10 to 65, n=11]; p=0.9; Figure 6) However, 3 veterinary journals displayed long intervals from acceptance to publication times: Australian Veterinary Journal, Journal of the American Veterinary Medical Association, and Journal of Veterinary Emergency and Critical Care. One of these is a society journal (Journal of the American Veterinary Medical Association), while the other two use a commercial publisher, commonly contracted by veterinary and medical societies (Wiley). For these 3 journals, the median acceptanceto-publication time was 265 days (IQR: 251 to 295) (Australian Veterinary Journal), 545 days (IQR: 499 to 603) (Journal of the American Veterinary Medical Association) and 572 days (IQR: 539 to 615) (Journal of Veterinary Emergency and Critical Care). Two additional journals (Veterinary and Comparative Orthopaedics and Traumatology, and Veterinary Surgery) had moderately long median intervals from acceptance to publication (137 and 148 days respectively). Only one medical or high-impact journal had an interval from acceptance to publication exceeding 100 days (British Journal of Surgery, 140 days).

**Figure 5.**
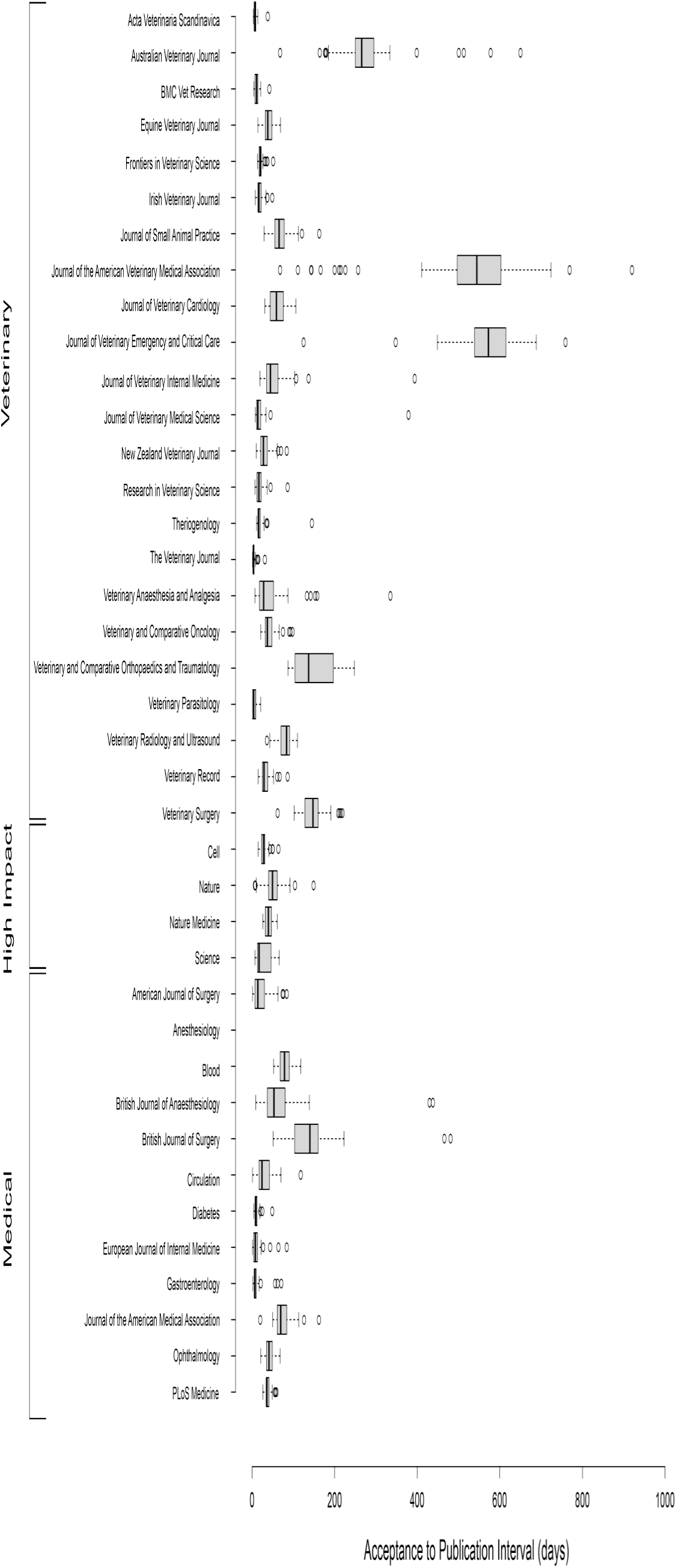
Box-and-whisker plots of the intervals from acceptance to first publication online of 50 articles for select veterinary, medical and high-impact journals. Whiskers denote the 5th and 95th percentiles. Circles denote articles outside the 5th and 95th percentiles.

**Figure 6.**
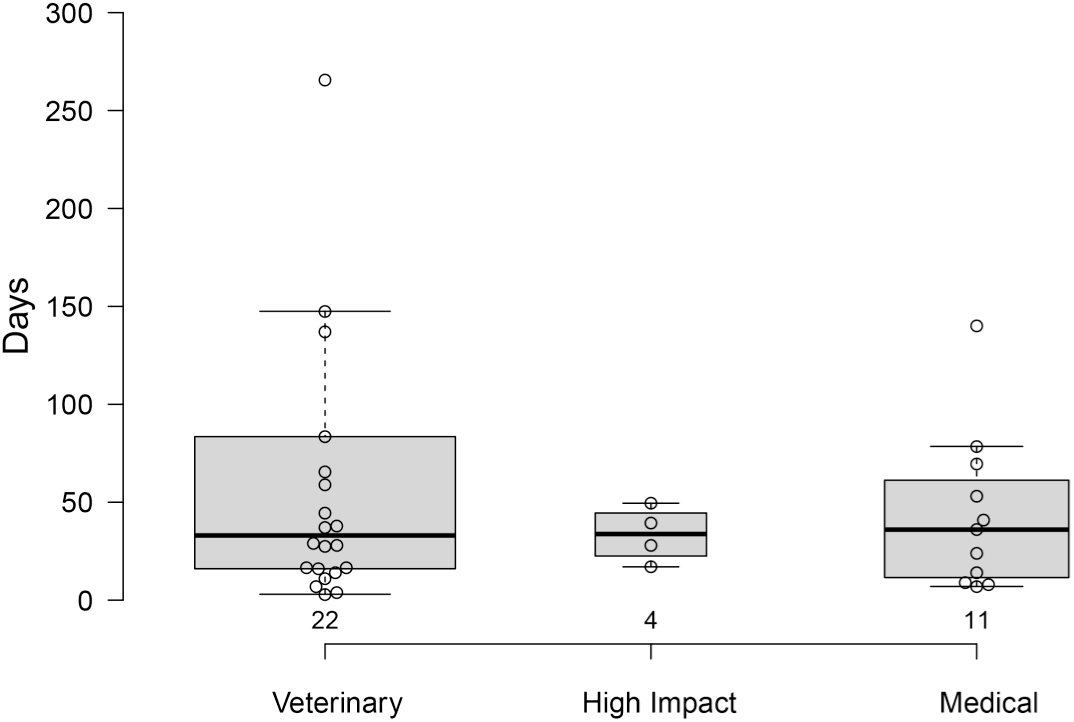
Box-and-whisker plots of median intervals from acceptance to publication in journals grouped by field (veterinary, medical, high impact). Groups did not differ from each other. Whiskers denote values within 1.5x the interquartile range. Circles denote individual values.

Intervals from acceptance to publication for journals published commercially (median: 36 days [IQR: 16 to 72, n=9]) did not differ from society-published journals (median: 33 days [IQR: 15 to 62, n=28]) (rho= -0.15, p=0.37, Figure 3B). The interval from acceptance to publication was associated with the total interval from submission to publication (rho= 0.7, p<0.001, Figure 4B).

### SOME VETERINARY JOURNALS HAVE LONG INTERVALS FROM SUBMISSION TO PUBLICATION

Figure 7 shows the intervals from submission to publication for all journals (Table 1). The total intervals, from submission to publication did not differ between fields (veterinary: 224 days [IQR: 171 to 288, n=23]; high-impact: 213 days [IQR: 190 to 288, n=4]; medical: 172 days [IQR: 146 to 198, n=9]; p=0.15; Figure 8). For most journals (28/34) the interval from submission to acceptance accounted for 70% to 90% of the total interval from submission to publication. However, for four journals, the intervals from submission to acceptance and from acceptance to publication were similar (approximately 50%), and for two of the three journals with the longest total interval from submission to publication, the interval from submission to acceptance accounted for 20% to 25% of the total interval from submission to publication. Specifically, the three veterinary journals with the longest intervals from acceptance to publication time displayed median intervals from submission to publication of 551 days (IQR: 429 to 761) (Australian Veterinary Journal), 665 days (IQR: 598 to 804) (Journal of the American Veterinary Medical Association) and 761 days (IQR: 714 to 822) (Journal of Veterinary Emergency and Critical Care) (Table 1). Because of their equally long intervals from submission to acceptance and acceptance to publication, Veterinary Surgery and the Journal of Comparative Orthopaedics and Traumatolology had total intervals from submission to publication exceeding 300 days. Only one other journal had a median interval from submission to publication approaching one year (Nature Medicine, 354 days); however, in contrast to the veterinary journals, this was caused by a long interval from submission to acceptance, rather than from acceptance to publication. Intervals from acceptance to publication for journals published commercially (median: 198 days [IQR: 174 to 248, n=8]) did not differ from society-published journals (median: 224 days [IQR: 165 to 270, n=27]) (p=0.7) and were not associated with impact factor (rho= -0.28, p=0.11) (Figure 3C).

**Figure 7.**
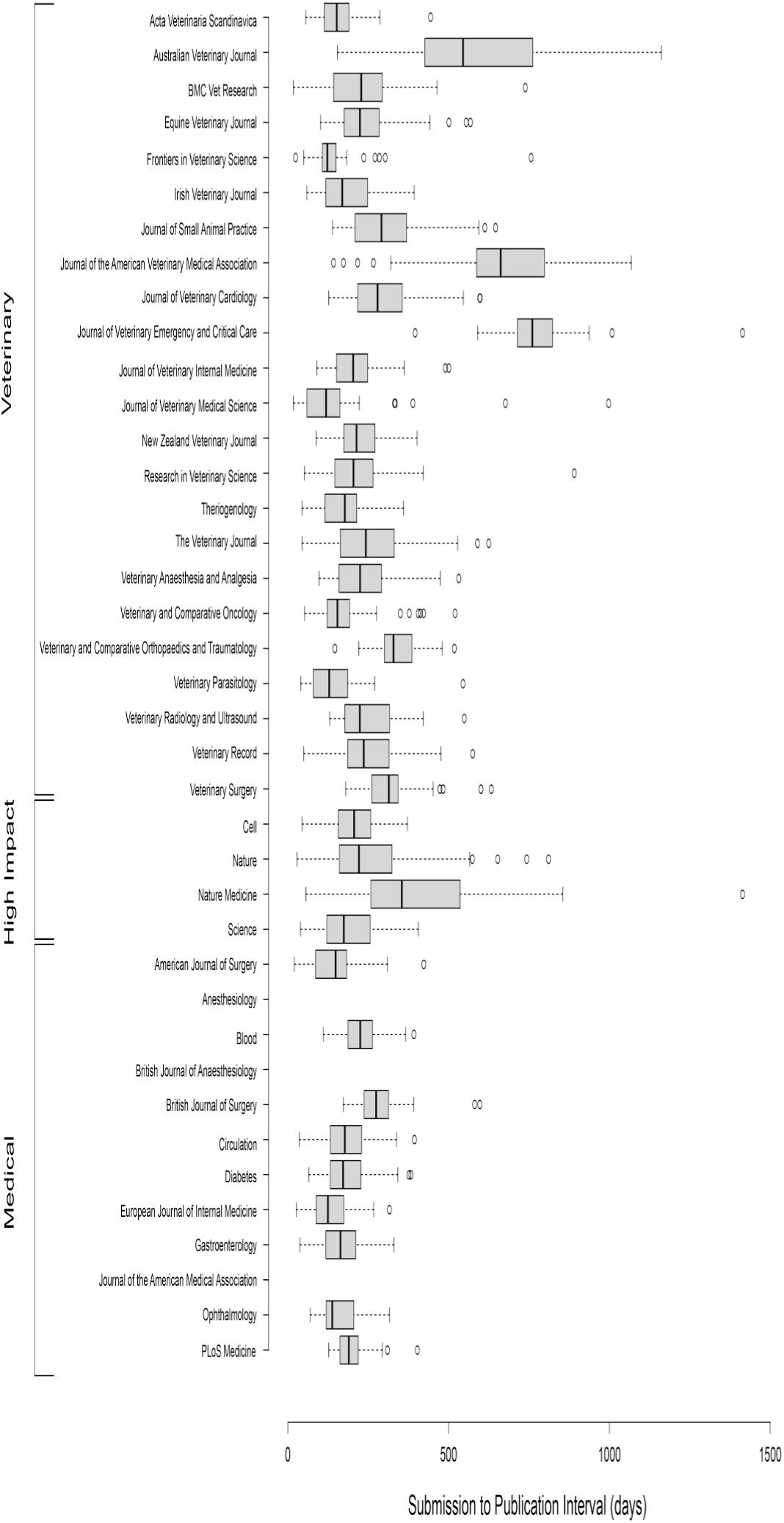
Box-and-whisker plots of time from submission to first publication online of 50 articles for select veterinary, medical and high-impact journals. Whiskers denote the 5th and 95th percentiles. Circles denote articles outside the 5th and 95th percentiles.

**Figure 8.**
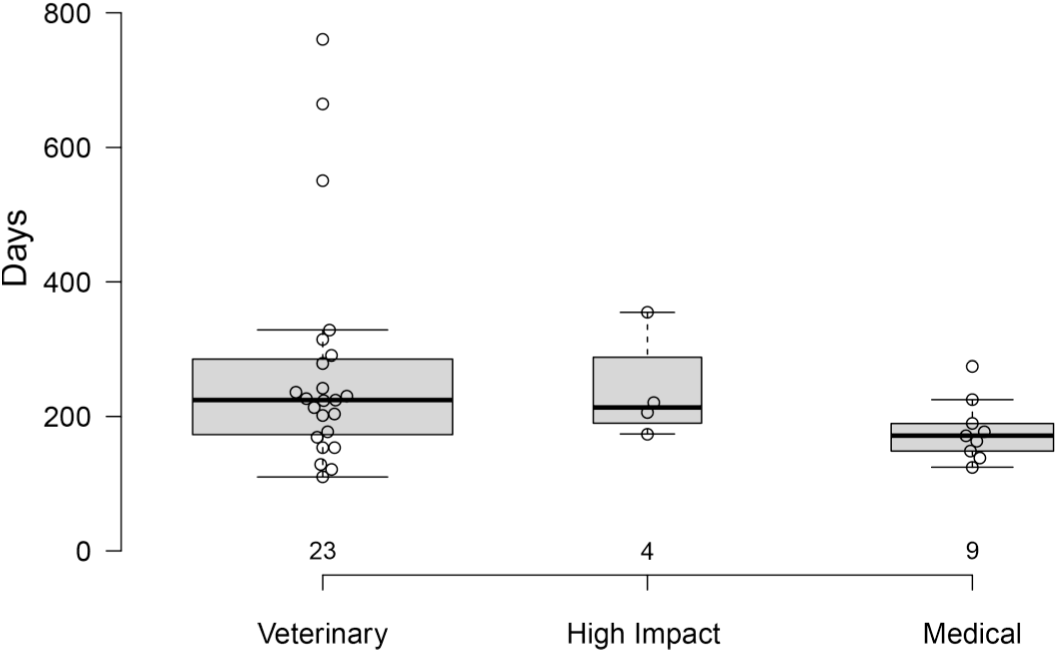
Box-and-whisker plots of median times from submission to publication in journals grouped by field (veterinary, medical, high impact). Groups did not differ from each other. Whiskers denote values within 1.5x the interquartile range. Circles denote individual values.

## DISCUSSION

Our study suggests that publication speed of veterinary research articles is similar to that of medical and general science high-impact journals, with a few notable exceptions. The interval from submission to acceptance appears to be similar across most journals, with relatively narrow interquartile ranges for most journals, and with most research manuscripts being accepted for publication within 9 months. Similarly, for most veterinary journals the interval from acceptance to online publication was similar that of medical and high-impact journals, with a few notable exceptions. Three veterinary journals had considerably longer median intervals between acceptance of a manuscript and its appearance online than most other journals we examined: The Australian Veterinary Journal, The Journal of the American Veterinary Medical Association and, somewhat ironically, the Journal of Veterinary Emergency and Critical Care. The last two of these took a median of 18 months to publish the information online after accepting the manuscripts. Therefore, our data suggest that the review process (the interval from submission to acceptance) is similar across veterinary clinical journals, and, furthermore, similar to that of medical or high-impact general science journals. Instead, the greatest impediment to prompt publication by some veterinary journals appears to lie with the time taken from accepting a manuscript to its appearance online.

Our data are similar to previous studies of publication speed for various medical journals. In most of those studies, median intervals from submission to acceptance times were approximately 5 to 6 months.^9^^,11–15^ A larger analysis by Daniel Himmelstein suggests that intervals from submission to acceptance has been reasonably static over the last 30 years, hovering around 120-140 days,^18^ with some variability between journals. The rigorous review expected from high-impact journals could plausibly lead to requests for additional experiments or studies after initial review and before final acceptance of a manuscript, thereby extending the interval from submission to acceptance; however, we did not find any such effects.

Our data show that most veterinary journals publish studies online soon after accepting the manuscript, some within a week. For example, Veterinary Parasitology, Acta Veterinaria Scandinavica and The Veterinary Journal all had intervals from acceptance to publication of less than a week. Many others had intervals less than one month. This is consistent with Himmelstein's data suggesting that intervals from acceptance to publication have dropped since 2000, and most dramatically since 2005, when online publication became increasingly available.^18^ His data suggest that median time to publication after acceptance is 25 days and has remained reasonably static over the last 10 years. However, in our study, three journals (Australian Veterinary Journal, Journal of the American Veterinary Medical Association, and Journal of Veterinary Emergency and Critical Care) had unusually long intervals between acceptance and online publication – 9 months for one journal and 18 months for the other two. Our evidence suggests that current publishing technology allows for rapid online publication by commercial publishers and society-published journals soon after acceptance (median time of 36 days). We did not ask the editors of these three journals to explain the unusually long time for online publication of accepted manuscripts and cannot speculate as to why such a long lag exists in these instances.

Evidence suggests that articles are much more likely to be cited if they are published online and freely available, consequently increasing the impact factor of the journal.^19^^,^^20^ However, similar to findings of some investigators,^7^^,^^12^^,^^15^ but contrary to other investigators,^11^ we found no association between intervals from acceptance to publication and impact factors. Furthermore, we found no association interval from acceptance to publication with field (veterinary, medical or high-impact) or type of publisher (society vs. commercial). Widely available digital technology should allow rapid publication regardless of who is publishing the journal or in what field the article is being published.

We found that both component intervals (submission to acceptance and acceptance to publication) were similarly and strongly associated with the total interval (from submission to publication) – in other words, journals that took longer to bring a submitted manuscript to print were slower to accept it and to publish it after acceptance.

We tried to obtain data for all the veterinary journals we considered relevant for clinical veterinary publications and to find comparable journals in the medical field. However, several journals, both veterinary and medical, fail to provide these data (Table 1), or provide only partial data. We do not know why journals choose to disclose some or none of this information, even when they are published by publishers who provide this information for other similar journals. This information should be available for authors to examine, so that they can make informed decisions about where to submit their manuscripts.

The 6-month median interval from submission to acceptance for scientific, peer-reviewed articles has remained static over 2 decades.^18^ The veterinary (and other) journals we studied show the same median duration for the interval from submission to acceptance – approximately 6 months. This interval is the sum of the time taken to find reviewers, review the manuscript, revise and re-submit the manuscript, and repeat these processes until the manuscript is finally accepted by the editors. None of these steps are easily shortened, nor would we expect them to differ between journals or fields, so we did not expect to find a relationship between the interval from submission to acceptance and the total interval from submission to publication. However, our data suggest that journals with longer total intervals from submission to publication also have longer intervals from submission to acceptance. Which step of this complex process is responsible for this relationship is hard to understand, given that reviewer behavior should be similar across journals, and authors should face similar delays in resubmitting manuscripts. Nevertheless, editors of journals with longer median intervals from submission to acceptance than most should examine where the delays to acceptance of a manuscript occur and determine if these factors are amenable to being addressed. We were encouraged to note that veterinary journals collectively have intervals from submission to acceptance similar to those of most other fields.

However, our data suggest that most journals could strive to decrease the intervals from submission to publication time by publishing manuscripts more quickly after acceptance. Several journals with intervals from acceptance to publication of less than a week demonstrate what is possible and provide a benchmark for other journals.

We examined only a “core” group of veterinary journals, which we considered “prominent” or “relevant” to veterinary clinical medicine. Additionally, we examined the publication metrics of only recently published articles. Finally, our comparator groups were small. Therefore, comparisons that did not detect a difference might suffer from a lack of power. However, our data agree with other published and unpublished data, appear robust from a visual inspection, and we would not anticipate that our findings would be invalidated by examining a larger cohort or with more data points per journal. If differences truly exist that we failed to detect, they would have to be inconsequential.

## CONCLUSION

In conclusion, our study shows that veterinary, medical and high-impact journals have similar publication speeds. Most journals have similar intervals from submission to acceptance and ranges. However, journals, regardless of field, discipline, publisher or impact factor, differ in the intervals from acceptance to publication, with several veterinary journals displaying long delays in bringing accepted manuscripts to readers.

## ACKNOWLEDGEMENTS

We thank the authors who responded to our request to provide publication information for articles published in the Journal of the American Veterinary Medical Association.

